# Allelic diversity of the pharmacogene *CYP2D6* in New Zealand Māori and Pacific peoples

**DOI:** 10.1101/2022.07.21.501043

**Authors:** Leonie M. Hitchman, Allamanda Faatoese, Tony R. Merriman, Allison L. Miller, Yusmiati Liau, Oscar E.E. Graham, Ping Siu Kee, John F. Pearson, Tony Fakahau, Vicky A. Cameron, Martin A. Kennedy, Simran D.S. Maggo

**Affiliations:** Department of Pathology and Biomedical Science, University of Otago, Christchurch, New Zealand; Christchurch Heart Institute, Department of Medicine, University of Otago, Christchurch, New Zealand; Biochemistry Department, University of Otago, Dunedin, New Zealand; Division of Clinical Immunology and Rheumatology, University of Alabama at Birmingham, Alabama, US; Auckland District Health Board, LabPLUS, Auckland City Hospital, Auckland, New Zealand; Pacific Trust Canterbury, Montreal Street, Christchurch, New Zealand; Department of Pathology, Center for Personalized Medicine, Children’s Hospital Los Angeles, Los Angeles, California, USA

**Keywords:** *CYP2D6*, pharmacogenetics, Polynesian, Māori, Pasifika, drug response, haplotype, ancestry

## Abstract

The enzyme cytochrome P450 2D6 (CYP2D6) metabolises approximately 25% of commonly prescribed drugs, including analgesics, anti-hypertensives, and anti-depressants, among many others. Genetic variation in drug metabolising genes can alter how an individual responds to prescribed drugs, including predisposing to adverse drug reactions. The majority of research on the *CYP2D6* gene has been carried out in European and East Asian populations, with Indigenous and minority populations greatly underrepresented. However, genetic variation is often population specific and analysis of diverse ethnic groups can reveal differences in alleles that may be of clinical significance. For this reason, we set out to examine the range and frequency of *CYP2D6* variants in a sample of 202 Māori and Pacific people living in Aotearoa (New Zealand). We carried out a long PCR to isolate the *CYP2D6* region before performing nanopore sequencing to identify all variants and alleles in these samples. We identified eleven novel variants, three of which were exonic missense variations. Six of these occurred in single samples and one was found in 19 samples (9.4% of the cohort). The remaining four novel variants were identified in two samples each. In addition, five new suballeles of *CYP2D6* were identified. One striking finding was that CYP2D6*71, an allele of unknown functional status which has been rarely observed in previous studies, occurs at a relatively high frequency (9.2%) within this cohort. These data will help to ensure that *CYP2D6* genetic analysis for pharmacogenetic purposes can be carried out accurately and effectively in this population group.

## Introduction

Pharmacogenetics is the study of genetic variants which impact on an individual’s response to drugs, with the aim of guiding prescription practices to improve healthcare quality and outcomes (Bank et al., 2018;Cacabelos et al., 2019). *CYP2D6* is one of the most studied pharmacogenes. This gene encodes an enzyme (CYP2D6) which is expressed in the liver and responsible for metabolising approximately 25% of commonly prescribed drugs or prodrugs (Nofziger et al., 2020).

The *CYP2D6* gene is highly polymorphic, with many recorded single nucleotide variants (SNVs), structural variants such as small insertions/deletions, larger copy number variants including whole gene deletions or duplications, as well as hybrid genes formed by recombination with the closely related pseudogene (*CYP2D7*) that is in close proximity (Nofziger et al., 2020). Over 150 *CYP2D6* alleles have so far been identified, each of which is allocated a name using a “star” nomenclature scheme, and tracked within the PharmVar database. Minor variations are often allocated a “suballele” designation (Gaedigk et al., 2020;Nofziger et al., 2020).

Variants of the gene may impact CYP2D6 function, ranging from completely inactivating the enzyme through to elevating its activity, although the impact of many variants is yet to be quantified. Where the functional impact of alleles is known or can be inferred, individuals can be categorised into one of four metaboliser phenotypes – ultrarapid metaboliser (UM), normal metaboliser (NM), intermediate metaboliser (IM), and poor metaboliser (PM), each of which describes activity of the CYP2D6 enzyme (Gaedigk et al., 2008). Depending on the type of medication, individuals defined as poor or ultrarapid metabolisers are at particular risk of experiencing drug toxicity or poor drug response (Cacabelos et al., 2019).

Interethnic differences in allele distribution or frequencies are evident for *CYP2D6* (Gaedigk et al., 2017;Zhou et al., 2017;Koopmans et al., 2021). For example, *CYP2D6*10* is reported with a frequency of 45% in East Asian populations, compared to only 1.6% in Europeans (Zhou et al., 2017). Furthermore, the PharmGKB database of *CYP2D6* variants (Whirl-Carrillo et al., 2021) includes data from over 64,000 Europeans compared to less than 800 individuals from the Oceanian biogeographic grouping, defined as pre-colonial populations of the Pacific, including Hawaii, Australia, New Zealand and Papua New Guinea (Huddart et al., 2019). Many of the existing Oceania’ studies focus on individuals from Papua New Guinea and Melanesia, which will not fully represent genetic diversity of Oceanian populations. Aotearoa (New Zealand) was the last major landmass to be inhabited, with Māori settlers arriving about 730 years before present (BP)(Gosling and Matisoo-Smith, 2018). Māori and Pacific Island populations have origins in South-East Asia, before migrations brought them to remote Oceania around 3000BP (Gosling and Matisoo-Smith, 2018). There have been very few studies specifically examining pharmacogenetic variability in Māori and Pacific Island people (Wanwimolruk et al., 1995;Wanwimolruk et al., 1998;Lea et al., 2008). To ensure equitable application of pharmacogenetic tests that detect clinically relevant alleles in people of all ancestries, it is important that such pharmacogenetic variants are identified and quantified by analysis of appropriate population samples.

The choice of technology employed for studies of interethnic diversity in *CYP2D6* is critical. Many prior studies have used targeted genotyping arrays or allele-specific polymerase chain reaction (PCR) methods, targeting only known alleles of interest, often derived from analysis of Europeans (Carvalho Henriques et al., 2021). These analyses are relatively cheap and straightforward to carry out, but unknown or rare alleles will not be detected (and potentially reported as *1 if not positive for another allele of interest). A more effective approach to identify all variants and determine haplotypes is to employ long-read, single molecule nanopore DNA sequencing on a PCR product encompassing the entire *CYP2D6* region (Liau et al., 2019). In this paper, we applied these methods to characterise the allelic landscape of *CYP2D6* in two cohorts of New Zealand Māori or Pacific Island people.

## Materials and Methods

### Study Cohorts

A total of 202 samples were drawn from two existing studies. Thirty-five of these samples were from a study called Genetics of Gout, Diabetes, and Kidney Disease in Aotearoa New Zealand, which recruited individuals aged ≥16 years primarily from the Auckland, Waikato and Christchurch regions of Aotearoa (New Zealand) (Krishnan et al., 2018). The remaining 167 samples were from the Pasifika Heart Study (Faatoese *et al*. manuscript in preparation). This study involved 200 Pacific participants aged 20 – 64 years, selected from the patient register of a Pacific-led primary healthcare clinic (Pacific Trust Canterbury, Christchurch, NZ), a low-cost health provider largely serving Pacific residents of Christchurch. Screening clinics for the Pasifika Heart Study were held from May 2015 – June 2016 at the Pacific Trust Health Clinic and the Nicholls Research Centre, University of Otago, Christchurch. In both studies, ethnicity was self-reported and the ethnicities of each participant’s four grandparents were also documented. The majority of samples were Polynesian, self-reporting as Samoan, Tongan, or Māori. Other included ethnicities were Cook Island Māori, Fijian (Melanesian), Tokelauan, Kiribati, and Niuean.

Ethical approval for the study Genetics of Gout, Diabetes, and Kidney Disease in Aotearoa New Zealand was given by the NZ Multi-Region Ethics Committee (MEC/05/10/130; MEC/10/09/092; MEC/11/04/036). The Pasifika Heart Study was approved by the New Zealand Health and Disability Ethics Committee (14/CEN/72/AM04). Participants in both studies gave written, informed consent.

**Table 1.** Ethnicity details of participants.

|  | Primary Ethnicity | Other Ethnicity 1 | Other Ethnicity 2 | Other Ethnicity 3 |
| --- | --- | --- | --- | --- |
| Samoan | 80 | 9 | 2 | 1 |
| Tongan | 53 | 3 |  |  |
| Fijian | 35 | 6 |  | 1 |
| NZ Māori | 23 | 1 | 1 |  |
| Cook Island Māori | 5 | 3 |  |  |
| NZ European | 3 | 11 | 1 |  |
| Kiribati | 1 |  |  |  |
| Tokelauan | 1 |  |  |  |
| Niuean | 1 |  |  |  |
| Grand Total | **202** | **33** | **4** | **2** |

### CYP2D6 Long PCR

*CYP2D6* was amplified by PCR as a 6.6kb product, including the whole gene as well as upstream and downstream non-coding regions. Duplication and deletion primers were used to identify the presence of a *CYP2D6* duplication or deletion via amplification of a secondary 3.5kb fragment if present (Gaedigk et al., 2007;Liau et al., 2019;Maggo et al., 2019b). Primer sequences are provided in Supplementary Table 1.

### Barcoding of PCR amplicons

Oxford Nanopore Technologies (ONT) PCR Barcoding Expansion 1-96 kit was used to add specific barcodes onto the *CYP2D6* amplicons. This allowed sample pooling in later steps. For a 50μL reaction, 0.5nM *CYP2D6* 6.6kb tailed amplicon was amplified in the presence of 1x LongAMP buffer, 0.3mM Kapa dNTPs, 5 units of LongAMP Hot Start *Taq* DNA polymerase, 0.2μM barcode and ultra-pure H_2_O to make up the final volume. This followed the ONT protocol SQK-LSK109, (version PBAC96_9069_v109_revO_14Aug2019). The PCR began with an incubation at 95°C for 3 minutes, followed by 15 cycles of 95°C for 15 seconds, 62°C for 15 seconds, 65°C for 7 minutes, and finally 65°C for 7 minutes, as recommended by ONT.

### Magnetic Bead Purification

Magnetic beads were used to purify the PCR products both prior to and following the barcoding step. This removed DNA fragments shorter than 3-4kb (to avoid off-target amplification) and other impurities. Magbio beads (MagBio Genomics Inc, Gaithersburg, Maryland, USA) were pelleted and re-dissolved in a buffer with 10mM Tris-HCl, 1.6M NaCl, 1mM EDTA pH 8, 11% (w/v) PEG 8000, 0.20% (v/v) Tween-20, and ultra-pure water to make a 10mL bead solution (Nagar and Schwessinger, 2018). DNA samples were purified as described previously (Liau et al., 2019).

### Nanopore Sequencing Library Preparation

DNA libraries of barcoded amplicons were prepared according to the 1D PCR barcoding (96) amplicons ONT protocol, SQK-LSK109. Briefly, after pooling the barcoded amplicons, about 200 fmoles of the DNA pool was subjected to end-repair and ONT adapter ligation. After purification, approximately 210 ng (~ 50 fmoles) of the library was introduced to the MinION flowcell (R9.4.1) and sequenced for up to 48 hours on the GridlON X5 nanopore sequencer (ONT, UK). Some sequencing was also completed on a FLO-FLG-001 flongle (R9.4.1) loaded with approximately 90 ng (~ 20 fmoles) of library.

### Nanopore Sequencing Data analysis

A previously designed pipeline was used to analyse the data generated (Graham et al., 2020). The GridION platform conducted real-time filtering, basecalling, and demultiplexing (separating and binning each sample per barcode) using Guppy version 5.0.12 (ONT, UK). A quality threshold for sequencing was set, specifying reads between 6 - 8kb in length and a Qscore of >9. An end-to-end pipeline for data management was designed in a conda environment using a snakemake workflow (Mölder et al., 2021). The process included utilising the output of the GridION to initially map the FASTQ files generated against a reference sequence (CYP2D6_NG008376.3) using MiniMap2 version 2.20-r1061 (Li, 2018). SAMtools (version 1.7) (Li et al., 2009) was employed to perform indexing at various stages of this pipeline. Nanopolish (version 0.13.2) (Quick et al., 2016) was used to analyse variant calls. Whatshap (version 0.17) (Martin et al., 2016) was used to phase the VCF files generated by Nanopolish. Variants were then matched to *CYP2D6* star alleles using the PharmVar database (Pharmacogene Variation Consortium) and unmatched variants were identified and marked as potentially novel to be taken forward for later validation. Finally, Stargazer v1.0.8 (Lee et al., 2019), an automated tool for pharmacogenetic star allele assignment, was used to confirm the *CYP2D6* star allele assigned to each sample. Star allele frequencies between studies were compared with two sided Fisher’s exact test and considered significant at p=0.05.

### Sanger sequencing

To validate potential novel alleles, Sanger sequencing was performed following a nested PCR on the relevant *CYP2D6* gene locations, using the long amplicon as a template as described (Maggo et al., 2019a; Maggo et al., 2019b). Primer sequences are provided in Supplementary Table 2.

## Results

From an initial 282 samples, 202 were successfully sequenced and analysed at all steps. Excluded samples included those that failed to amplify the 6.6kb *CYP2D6* PCR product (n=52), or samples with a low read depth (less than 50 reads; n=28). Genomic DNA samples which failed to amplify were checked on a TapeStation platform (Agilent, Santa Clara, USA) and were usually found to be fragmented, with this fragmentation being the likely cause of long-PCR failure. 120 of the samples were sequenced with an R9.4.1 flow cell, and the remaining 82 were sequenced with a FLG-001 Flongle, with both flow cell types run on the GridION X5.

### CYP2D6 Genotyping Results

Nine different *CYP2D6* star alleles were identified amongst 202 individuals, including the *5 gene deletion (Table 2). No *CYP2D6* gene duplications were observed. The most common allele was *1 with an allele frequency of 0.48. Following this, in order of frequency, were *10, *2, and *71. Four of the identified star alleles, *4, *5, *10, and *41, are known to decrease or completely inactivate the function of the CYP2D6 liver enzyme, while *1, *2, and *35 have no impact. *43 and *71 have unknown functional effects on CYP2D6.

**Table 2.** Star allele frequencies and associated information.

| Star allele | Allele frequency<br>(n=202) | Number of haplotypes observed<br>(n=404) | Activity score<sup>1</sup> | Impact |
| --- | --- | --- | --- | --- |
| *1 | 0.478 | 193 | 1 | Normal function |
| *2 | 0.141 | 57 | 1 | Normal function |
| *4 | 0.017 | 7 | 0 | No function |
| *5 | 0.012 | 5 | 0 | No function |
| *10 | 0.183 | 74 | 0.25 | Decreased function |
| *35 | 0.007 | 3 | 1 | Normal function |
| *41 | 0.067 | 27 | 0.5 | Decreased function |
| *43 | 0.002 | 1 | ? | Uncertain function |
| *71 | 0.092 | 37 | ? | Uncertain function |

<sup>1</sup> Activity score and associated metaboliser phenotype as defined by Caudle et al (2020).

Stargazer 1.0.8 was used to call star alleles for the first 120 samples, using variants aligned to hg19. Of these 120, eleven were called incorrectly. Five of the incorrect haplotype assignments were due to the presence of a *5 allele, while the remaining six carried *71 variants.

### Predicted Metaboliser Phenotypes

The metaboliser status was inferred from the genotype of each sample using the diplotype activity score as previously defined (Caudle et al., 2020). In this classification, diplotypes with an activity score of 0 are classified as poor metabolisers, 0.25-1 are intermediate metabolisers, 1.25-2 are normal metabolisers, and scores above 2 are ultrarapid metabolisers. Using this definition, the majority (74%) of individuals within the cohort were classified as normal metabolisers. No individuals were identified as being ultrarapid or poor metabolisers, and a small subset (8%) of our study cohort were intermediate metabolisers. Almost 20% of the characterised samples had a metaboliser status that could not be assigned due to the presence of one or more *71 or *43 alleles (Table 3).

**Table 3.** Inferred metaboliser frequencies within cohort.

| Inferred Phenotype (from genotype) | Number | Frequency |
| --- | --- | --- |
| Ultrarapid | 0 | 0 |
| Normal | 150 | 0.743 |
| Intermediate | 16 | 0.079 |
| Poor | 0 | 0 |
| Unknown | 36 | 0.178 |

### Novel Variants

We identified eleven potentially novel variants and validated them using Sanger sequencing (Table 4). Seven variants occur within an intron of the gene, and four are exonic variants, three of which are non-synonymous. These have the potential to form new star alleles or new suballeles.

**Table 4.** Novel Variants.

| Variant position<br>(CYP2D6 reference)<sup>1</sup> | SNP rsID | Type of variant | Number of times<br>observed | Allelic designation |
| --- | --- | --- | --- | --- |
| 5252 (Exon 2) | rs/na | Missense A>G (Gln > Arg) | 1 | *new |
| 5709 (Intron 2) | rs369508051 | G>A | 19<sup>2</sup> | *41.new |
| 6128 (Exon 4) | rs755518310 | Missense G>A (Gly > Glu) | 1 | *71.new |
| 6239 (Intron 4) | rs564994275 | A>C | 1 | *4.new |
| 6242 (Intron 4) | rs778008161 | G>C | 1 | *1.new |
| 6419 (Intron 4) | rs/na | G>C | 2 | *1.new |
| 7557 (Intron 7) | rs376909251 | G>A | 1 | *1.new |
| 7574 (Intron 7) | rs/na | Indel T>TCAGCAC | 2 | *1.new |
| 8030 (Exon 8) | rs1602566413 | Missense G>C (Val > Leu) | 2<sup>2</sup> | *new |
| 8068 (Exon 8) | rs1037093492 | Synonymous C>T (Leu > Leu) | 2<sup>2</sup> | *2.new |
| 8202 (Intron 8) | rs769045995 | A>G | 1 | *41.new |

<sup>1</sup>Reference sequence AY545216 (1=sequence start) NG\_008376.3
<sup>2</sup>Variant also identified by Charnaud et al. (2022)

Six of the eleven novel variants were singletons (detected in only one individual each), four were identified in two individuals each, while a variant at position 5709 was found in nineteen people. In addition, five potentially novel suballeles were also identified (data not shown). These variants occur within the PharmVar CYP2D6 database but as part of different star alleles to those observed in this study. The majority exist on the haplotype without any core variants belonging to star alleles, so are by default *1. Two variants at position 8595 and 8645 are likely a new *10 suballele as all samples have the *10 core variants.

## Discussion

Of the nine broad ethnic groups defined by Huddart *et al*., and reported by PharmGKB, Oceania is by far the least represented (PharmGKB) (Huddart et al., 2019). We have successfully used nanopore sequencing to analyse the entire *CYP2D6* gene (including important upstream and downstream regions) of 202 Māori and Pacific Island individuals, coupled with analyses to call the star allele diplotypes of each participant. Nine star alleles were identified amongst this cohort, with frequencies ranging from 0.48 for *1, to 0.002 for *43. CYP2D6*71, a previously identified rare allele with unknown function, was identified at a relatively high frequency within this cohort, with an allele frequency of 0.092. From each individual’s diplotype, we were able to categorise the majority of the cohort into metaboliser phenotypes. We did not find any ultrarapid or poor metabolisers. A large percentage of participants (74%) were normal metabolisers and 8% had a predicted activity score within the range of intermediate metabolisers (Caudle et al., 2020). The remaining 18% were unable to be assigned to a metaboliser status due to carrying one or more copies of the *43 or *71 allele.

Eleven novel variants were also discovered, four of which are exonic. Five known variants were found within star alleles in which they have not been reported previously, potentially forming new suballeles.

We did not identify any poor metabolisers in the present study, as all non-functional alleles were found in diplotypes with a fully functional or partially functional allele. Non-functional alleles (*4 and *5 gene deletion) were present in ~3% of the cohort, so they do exist at a low frequency within the population and therefore could give rise to poor metabolisers phenotypes. In this regard, it is interesting to note that a pharmacological study using the probe drug debrisoquine in 101 Māori participants identified 5% as poor metabolisers (Wanwimolruk et al., 1995), and a similar study in 100 Polynesian participants found none were poor metabolisers (Wanwimolruk et al., 1998). Therefore, our genetic findings are reasonably congruent with these earlier studies.

However, *CYP2D6* allele frequencies observed in our study do not match allele frequencies previously reported by PharmGKB for the Oceanian region (PharmGKB). This is almost certainly because the Oceanian definition encompasses populations with diverse, complex migration and ancestral histories (Gosling and Matisoo-Smith, 2018). The Oceanian *CYP2D6* data currently available comes from studies that in total include less than 800 individuals. The largest study (Gutiérrez Rico et al., 2020) focused on 278 Ni-Vanuatu (Micronesian) subjects, and found the majority of individuals to carry at least one *1 allele (allele frequency of 0.804), significantly greater than our identified frequency of 0.478 (P = 4.2 x 10^-26^). In total, they identified nine different star alleles amongst their cohort, the majority of which had allelic frequencies of <0.01. *2 and *10 existed above this frequency (0.068 and 0.029, respectively) as well as a duplication of *1 at 0.025 (Gutiérrez Rico et al., 2020). Our study found no duplications and only two alleles had observed frequencies below 0.01 (*35, *43). Prior reports describing high rates of *CYP2D6* duplications in Oceania have been entirely based on Papua New Guinea samples (Sistonen et al., 2007;von Ahsen et al., 2010).

The relatively high frequency of the *71 allele we observed is one of the most striking findings from this study, for several reasons. Because *71 has thus far been considered a rare allele, it would not be routinely included in most *CYP2D6* genotyping panels. The functional impact of *71 is unknown, so we were unable to predict metaboliser status for 18% of our cohort. The first study to identify *71 was performed in a Han Chinese population (Zhou et al., 2009). This group suggested the allele could be non-functional due to the p.Gly42 >Glu substitution in the protein N-terminus, a region which is involved in membrane anchoring of the enzyme (Zhou et al., 2009). However, this is speculative and has yet to be tested. Other reports, in Micronesian populations, have described allele frequencies of 0.009 in a Ni-Vanuatu sample (Gutiérrez Rico et al., 2020), significantly less than our 0.092 (P = 3.9×10^-10^) and 0.058 in a large sample of Solomon Islanders (Charnaud et al., 2022), comparable but significantly less than our 0.092 (P=0.038).

## Conclusion

In conclusion, we have used a nanopore sequencing approach to identify *CYP2D6* star alleles and respective frequencies from 202 individuals of Māori or Pasifika ethnicity. We identified nine known star alleles in this cohort, including CYP2D6*71, a previously identified rare allele of unknown function, which comprised over 9% of the alleles detected. The frequency of this allele prevented us from inferring metabolic function for nearly 20% of this cohort, and clearly, understanding the functional impact of *71 using clinical phenotyping approaches or *in vitro* analyses is a priority area for future research. Should this allele have an altered function, it would be important to ensure it is included in pharmacogenetic testing panels, particularly if to be used in the Oceania region.

Our analysis did not reveal any *CYP2D6* duplications in this cohort, and overall our findings highlight the significant limitations of the Oceania biogeographic grouping for pharmacogenetic research, given the wide allele frequency differences we observe relative to studies on other Oceanian ancestral groups.

Finally, this study has further demonstrated the utility of nanopore sequencing for highly variable genes like *CYP2D6*, with long-read sequencing providing the advantage of analysing the entire gene, including intronic regions, at high throughput and relatively low cost. It is to be hoped this research will contribute to a more equitable uptake of *CYP2D6* pharmacogenetics in people of Māori and Pacific ancestry.

---

**Supplementary Table 1.**
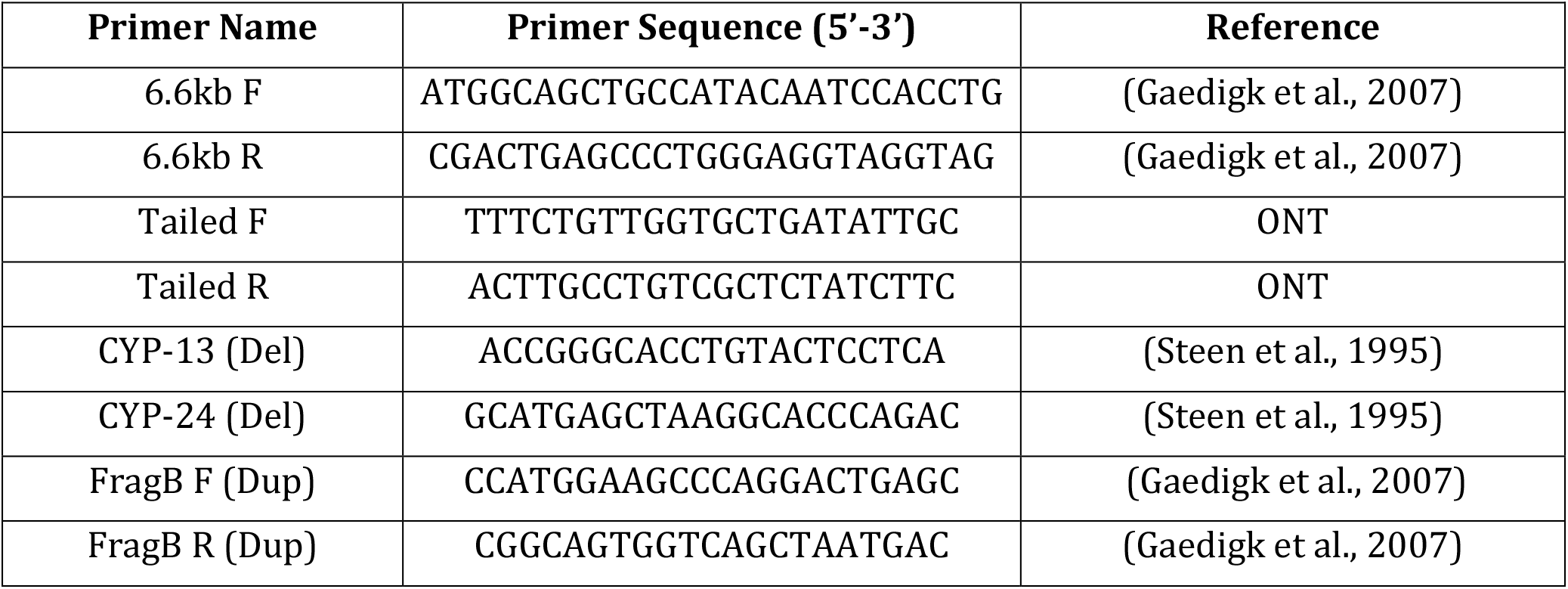
Primer Sequences for CYP2D6 amplification.

**Supplementary Table 2.**
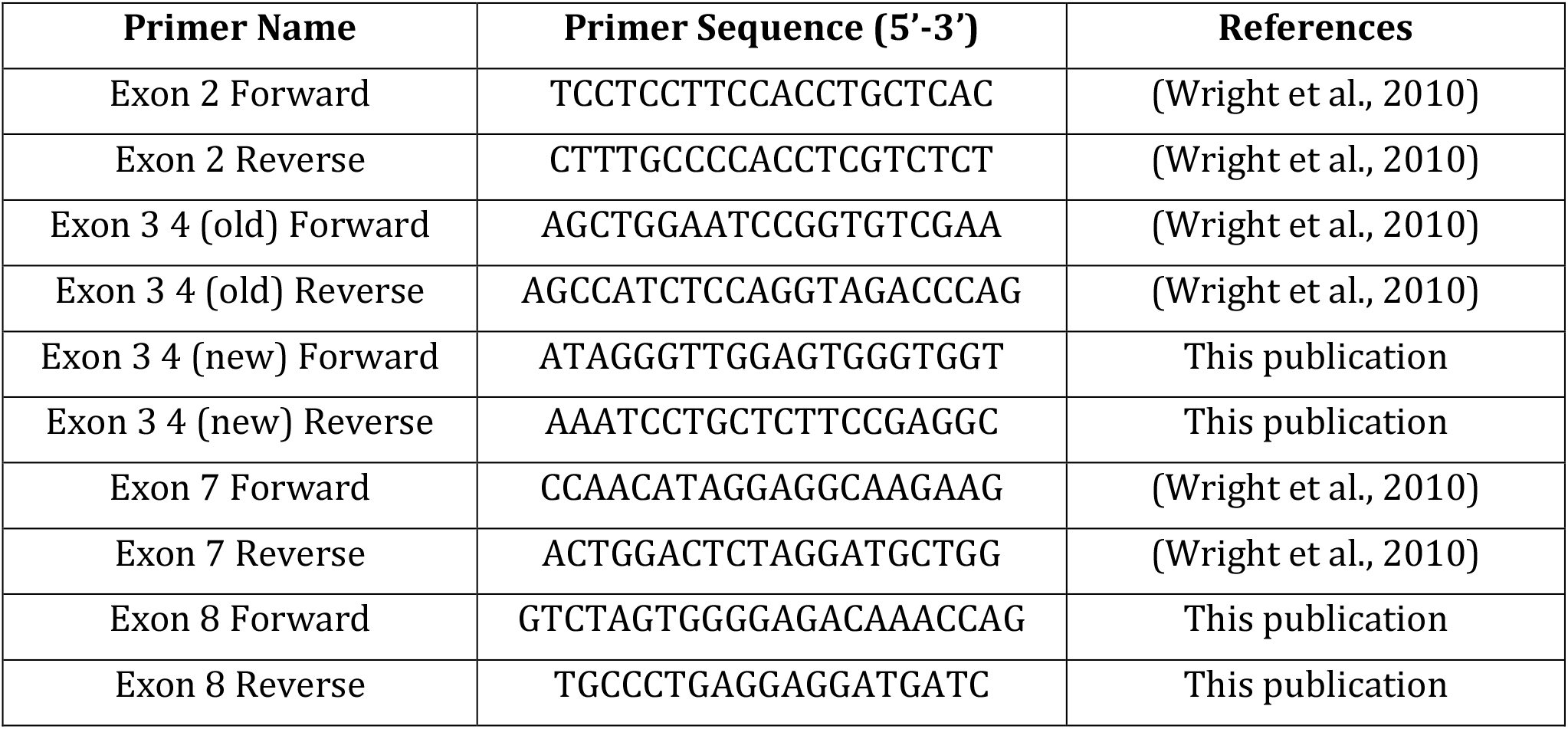
Primer Sequences for novel variant confirmation.

